# Deuteration provides a general strategy to enhance azobenzene-based photopharmacology

**DOI:** 10.1101/2023.11.09.566420

**Authors:** Kilian Roßmann, Alberto J. Gonzalez-Hernandez, Rahul Bhuyan, Karl Börjesson, Joshua Levitz, Johannes Broichhagen

## Abstract

Herein, we present deuterated azobenzene photoswitches as a general means of enhancing photopharmacological molecules. Deuteration can improve azobenzene performance in terms of light sensitivity, photoswitch efficiency, and photoswitch kinetics with minimal alteration to the underlying structure of the photopharmacological ligand. We report synthesized deuterated azobenzene-based ligands for the optimized optical control of ion channel and G protein-coupled receptor function in live cells, setting the stage for the straightforward, widespread adoption of this approach.

## MAIN TEXT

Photopharmacology represents a powerful means of optically controlling biological function through the use of light-sensitive compounds^1,2^. In addition to its use in a wide variety of basic science applications^3–12^, photopharmacology has now entered clinical trials via KIO-301, a photoswitchable ion channel blocker with great potential for vision restoration^13^. This compound utilizes an azobenzene-based photoswitch, which represents one of the primary chemical moieties used in photopharmacological probes^14^. Despite their many advantageous properties^15^, azobenzene photoswitch performance is typically limited by light sensitivity, photoisomerization efficiency, and photoswitching speed which together reduce their ability to enable robust and rapid light-dependent control in complex biological systems. In recent years, multiple strategies have emerged to improve the properties of azobenzene-based photoswitches. Most typical has been derivatization of the azobenzene itself which can enhance critical photophysical properties (extinction coefficient, wavelength tuning, bistability etc.) by a diverse array of chemical modifications, introducing heterocycles or halogen atoms on the aromatic units, creating push-pull systems or the installment of sensitive “antennas” for 2-photon activation^14,16–25^. To enable improved target selectivity and genetic precision, covalent tethering to cysteines or self-labelling enzymes (e.g. SNAP-tag) has also been used. We recently reported a strategy to effectively improve tethered photopharmacology efficiency by branching multiple azobenzene switches onto the same molecule^26,27^.

Despite their utility, all of the aforementioned techniques involve chemical modifications to the underlying compound, thus altering the core structure of the molecule. This makes the process of improving photopharmacological ligands laborious and molecule-specific and raises the possibility that chemical modifications will not be tolerated due to constraints of the target molecule’s binding site. Methods that can be broadly applied to any azobenzene-based system without the need for compound-specific engineering are, thus, needed. One such option is to pursue deuteration, which introduces isotope effects without altering the structure of the chromophore itself. Motivated by recent studies showing that installation of deuterium can enhance fluorophore performance^28–30^ (**Figure 1A**), we asked if similar improvements may be obtained with azobenzene photoswitch chromophores (**Figure 1B**). Deuterated azobenzenes have been described before to investigate drug metabolism^31^, to study ^13^C shifts in NMR spectroscopy^32^, or to obtain “IR clean” switches^33^. However, to our knowledge, no reports exist that explore deuterated azobenzenes in photopharmacological settings.

**Figure 1:**
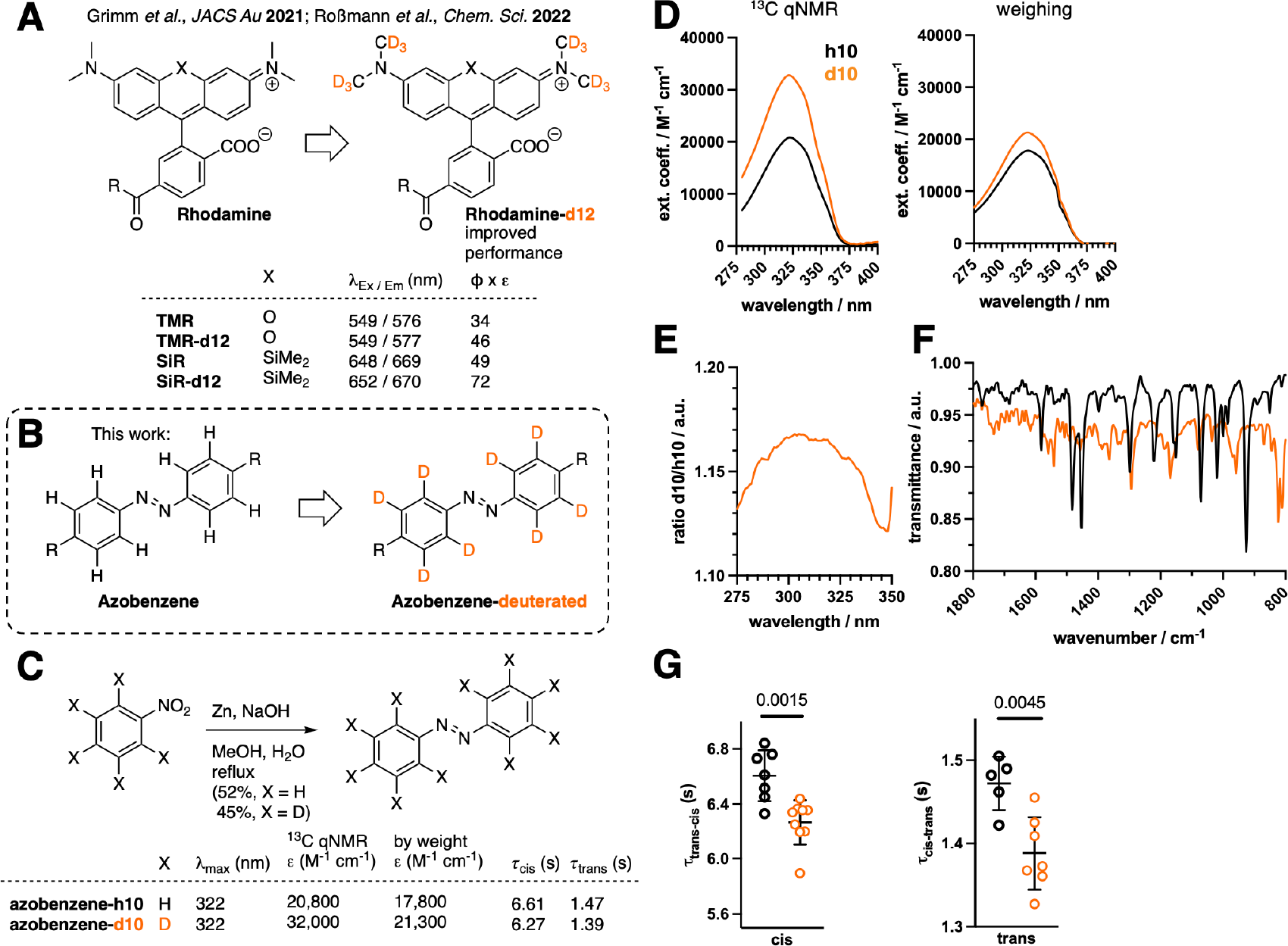
Deuteration strategy to enhance photophysical properties. **A)** Enhanced fluorophore performance was previously obtained by installing deuterated *N*-methyl rhodamines. While excitation/emission spectra show minute changes, brightness (Φ x ε) is drastically enhanced by deuteration. **B)** In this work, we extend this concept by perdeuterating azobenzene chromophores. **C**) Synthesis of azobenzene-d10 by reductive dimerization of nitrobenzene using zinc. Extinction coefficient and switching kinetics are increased by deuteration. **D-E**) UV/Vis spectra and extinction coefficient determination of azobenzene-h10 and azobenzene-d10 in DMSO by ^13^C qNMR (left) or by weighing (right). **E**) Ratio of azobenzene-d10 and azobenzene-h10 shows different absorbance characteristics. **F**) IR spectra of azobenzene-d10 and azobenzene-h10 shows distinct vibrational states. **G**) Switching kinetics for trans-to-cis (left) and cis-to-trans (right) photoconversion of azobenzene-h10 and azobenzene-d10 in DMSO. P-values from unpaired t-tests are reported in panel G.

We first aimed for the simplest model system, a ‘naked’ azobenzene with either 10 hydrogens (“AB-h10”) or 10 deuteriums (“AB-d10”) (**Figure 1B**, R = D). Synthesis was straightforwardly achieved from non-deuterated or deuterated nitrobenzene using zinc as a reducing agent in refluxing methanol, and the desired azobenzenes were obtained in 51% and 45% yields, respectively (**Figure 1C**). We characterized the photophysical properties of these molecules and found that the maximal absorbance wavelength in DMSO remained unchanged at 322 nm (**Figure 1D**). Since no hydrogen atoms are present in azobenzene-d10, we next performed hydrogen-coupled, quantitative ^13^C NMR to determine concentrations using DMF as an internal standard, and found using UV/Vis spectroscopy that extinction coefficient was increased by >50% (ε_322 nm_= 20,800 versus 32,000 M^-1^ cm^-1^) for AB-d10 (**Figure 1D**). Impressed by this change, and to exclude distortions by nuclear Overhauser effects due to different nuclei, we confirmed this trend by weighing each compound and observed the extinction coefficient to be increased by ∼20% due to deuteration (ε_322 nm_= 17,800 versus 21,300 M^-1^ cm^-1^) (**Figure 1D**). We note that in both cases, AB-h10 was close to reported literature values of 22,400 M^-1^ cm-^1^ at 319 nm in methanol.^34^ Interestingly, by plotting the absorbance of the deuterated divided by the absorbance of the non-deuterated azobenzene, we observed a subtle change in spectra around the maximal absorbance peak (**Figure 1E**), indicating different vibrational states for the two molecules. To further examine this, we recorded IR spectra of AB-h10 and AB-d10 and found, as expected, differences in the fingerprint region (**Figure 1F**). Most relevant to potential photopharmacological applications, we also observed a clear acceleration in *trans*-to-*cis* (τ = 6.61 vs. 6.27 sec) and *cis*-to-*trans* (τ = 1.47 vs. 1.39 sec) photoswitching kinetics for azobenzene-d10 in DMSO (**Figure 1G**).

Encouraged by the above results indicating that deuteration can improve azobenzenes, we pursued a water soluble azobenzene-based compound since organic solvent effects do not recapitulate the cellular environment where most photoswitches are, ultimately, applied. We chose azobenzene quaternary ammonium (AQ) as a scaffold as it is bis-amidated and, thus, carries a charge for excellent water solubility. AQ has been used with various substituents to optically control potassium channels in a plethora of studies on nociception, vision restoration, and neuromodulation (**Figure 2A**)^13,23,35–39^. We synthesized deuterated AQ-d8 by oxidatively dimerizing phenylene diamine-d4 (**1**) with Dess-Martin periodinane to obtain a perdeuterated 4,4’-bisamine azobenzene **2**, before HBTU-mediated coupling to betaine and subsequent acylation using acetyl chloride (**Figure 2B**). We profiled AQ-h8 and AQ-d8 and found relatively unchanged maximal absorbance at 363 nm and 360 nm (**Figure 2C**), respectively. We determined extinction coefficients in water to be 15,200 M^-1^ cm^-1^ for both compounds via ^1^H qNMR using DMF as an internal standard (**Figure 2C**). We probed the change in UV/Vis absorbance by looking at the ratio of values for AQ-h8 and AQ-d8 and found subtle changes around the maximal absorbance value (**Figure 2D**). IR spectra also showed distinct shifts in vibrational motions (**Figure 2E**), indicating differences due to the deuterium isotopes. ^1^H qNMR measurements allowed us to determine photostationary states (as described previously^23^) in D_2_O under 385 nm, 500 nm and 525 nm irradiation where similar values were seen for both compounds (**Figure 2F**). We also determined quantum yields for *trans*-to-*cis* switching and found these to be similar (Φ(AQ-h8) = 32%; Φ (AQ-d8) = 31%)^40^. While such modest differences would likely not strongly influence performance in a photopharmacological setting, we observed that switching kinetics were much faster for AQ-d8 than AQ-h8 (*trans*-to-*cis*: τ = 9.91 vs. 5.42 sec; *cis*-to-*trans*: τ = 6.11 vs. 4.18 sec) (**Figure 2G**). Encouraged by this, we tested the ability of AQ-h8 and AQ-d8 to control the activity of large conductance voltage and calcium-gated (BK) potassium channels *via* patch-clamp electrophysiology in HEK 293 cells. We delivered 1 mM of AQ-h8 or AQ-d8 to the cytosol *via* the patch pipette and observed robust, reversible photo-block and photo-unblock by illuminating successively with 525 nm and 385 nm light (**Figure 2H**). While the efficiency of photoblock was similar for AQ-h8 and AQ-d8 (**Figure S1A, B**), AQ-d8 showed substantially faster photoswitch kinetics (**Figure 2I**). Importantly, given that AQ acts as a simple pore blocker, photocurrent kinetics likely serve as a direct readout of cis/trans switching kinetics.

**Figure 2:**
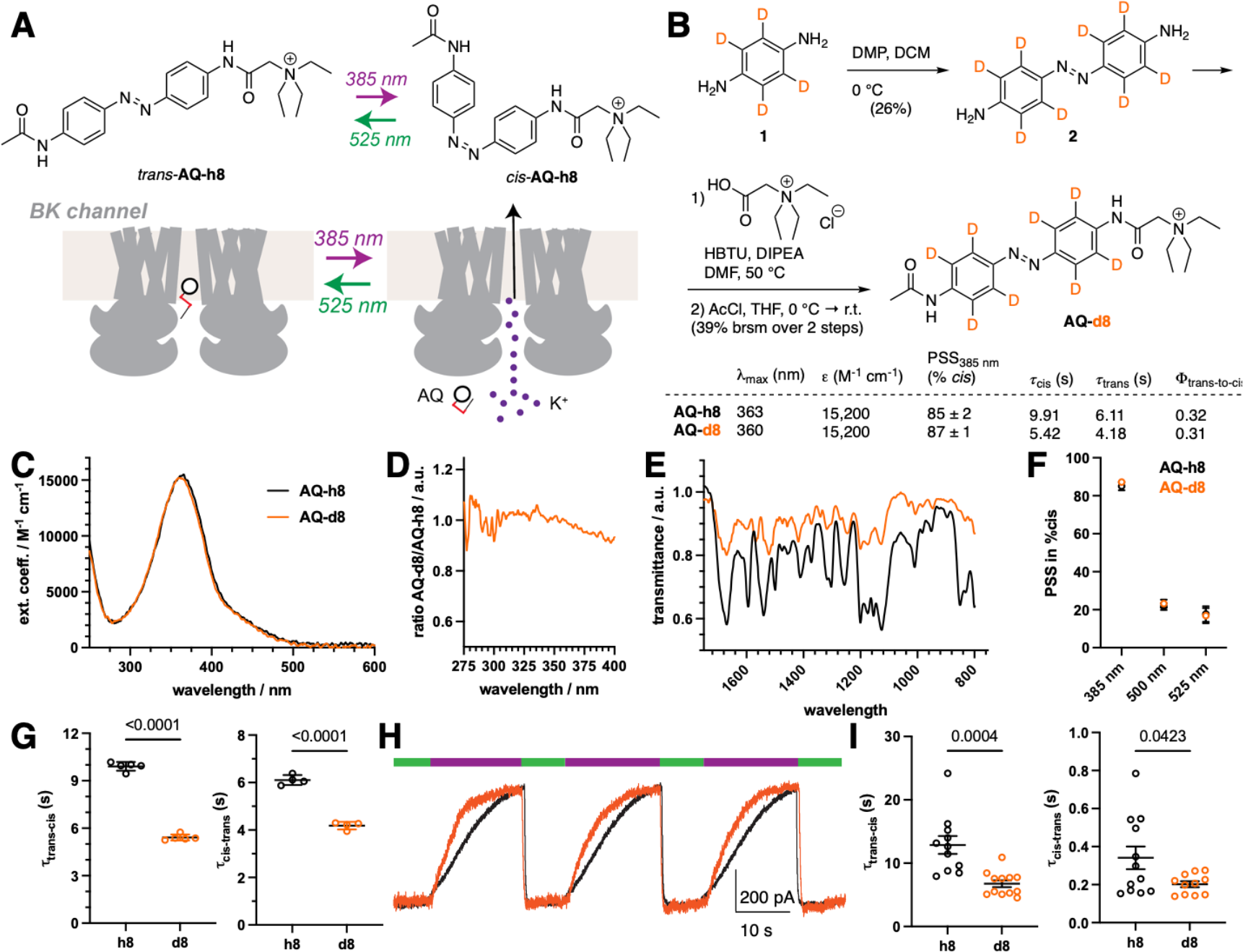
Deuteration enhances performance of a soluble, photoswitchable potassium channel blocker. **A)** *trans*-AQ blocks potassium channels at an intracellular site and unblocking can be achieved optically by applying 385 nm light, with reversibility using 525 nm light. **B**) Synthesis of AQ-d8 and summary of photophysical properties. **C**) UV/Vis spectra and extinction coefficient determination of AQ-h8 and AQ-d8 in water by ^1^H qNMR. **D**) Ratio of AQ-h8 and AQ-d8 shows different absorbance characteristics. **E**) IR spectra of AQ-h8 and AQ-d8 shows different vibrational states. **F**) Photostationary state occupancies under different wavelengths as assessed by ^1^H qNMR. **G**) *in vitro* Switching kinetics of AQ-h8 and AQ-d8. **H**) BK channel representative trace at +60 mV in response to 385 nm (purple) and 525 nm (green) light in the presence of AQ-h8 or AQ-d8. I) Quantification of channel photo-unblock (left) and photo-block (right) kinetics. P-values from unpaired t-tests are reported in panels G and I.

Photopharmacology can be merged with the power of genetic engineering by tethering photoswitchable ligands to a self-labelling tag (e.g. SNAP) on a protein of interest^26^. This approach yields excellent target selectivity due to the biorthogonal nature of labelling and rapid photoswitching kinetics due to the lack of ligand diffusion. We pioneered this approach by conjugating the SNAP-tagged metabotropic glutamate receptor 2 (SNAP-mGluR2), a neuromodulatory GPCR, with a “photoswitchable orthogonal remotely-tethered ligand” (PORTL) which enables rapid, reversible optical control of mGluR2 activity *ex vivo* and *in vivo* (**Figure 3A**)^23,27,41^. The PORTL ligand consists of a benzylguanine-azobenzene-glutamate (“BGAG”) photoswitch, with BGAG_12_-v2-h8 serving as a testbed for our deuteration strategy. Employing a previously described synthetic route (**Figure 3B; Supplementary Information**), we obtained deuterated BGAG_12_-v2-d8, which showed the same maximal absorbance wavelength of BGAG_12_-v2-h8 (**Figure 3B**; **Figure S2A**). BGAG_12_-v2-d8 labelled SNAP-mGluR2 transfected HEK293 cells with the same efficiency as its non-deuterated counterpart (**Figure S2B, C**). Using patch-clamp electrophysiology with G protein-coupled inward-rectifying potassium (GIRK) channels as a reporter, we observed robust and reversible responses by applying 385 nm (ON) and 525 nm (OFF) light (**Figure 3C**). When comparing photocurrents to the response to a saturating concentration of glutamate (1 mM), a clear increase in photoswitching efficiency was observed from ∼50% to ∼68% for BGAG_12_-v2-d8 (**Figure 3D**). In addition, we observed a faster ON response when 385 nm light was applied (**Figure 3E**). It should be noted that OFF kinetics, which do not change between BGAG_12_-v2-d8 and BGAG_12_-v2, do not recapitulate photoswitch kinetics in this system but are limited by biological signal termination processes. Nevertheless, the PORTL system allows for a clean readout since photoswitch concentration is determined by receptor expression level.

**Figure 3:**
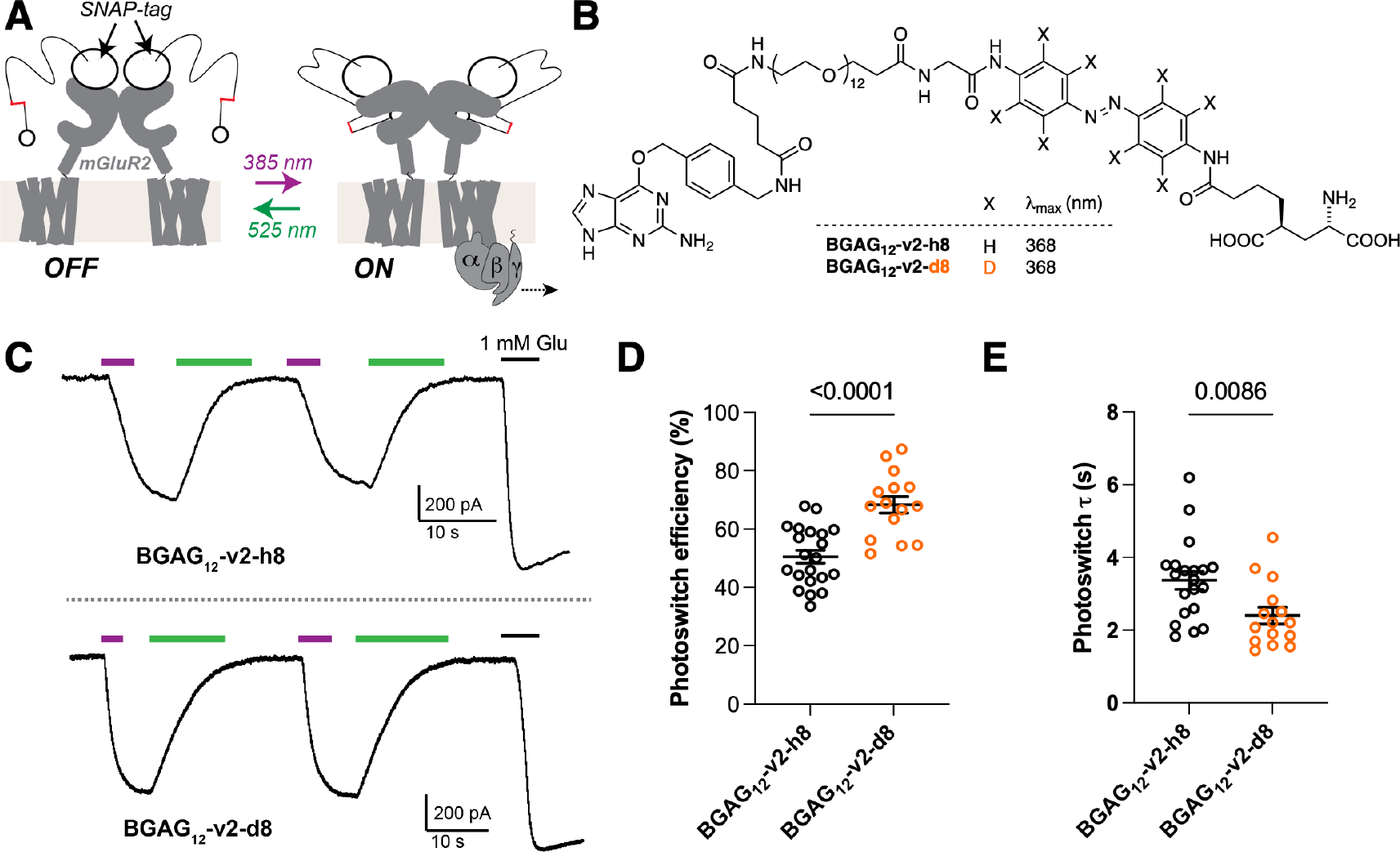
Deuteration enables more efficient, faster optical control of a GPCR via a tethered photoswitch. **A**) Schematic showing optical control of mGluR2-mediated G protein signalling via the PORTL “BGAG”. BGAG is attached to an N-terminal fused SNAP-tag and activates mGluR2 upon 385 nm light and can be turned off using 525 nm light. **B**) Structure of BGAG_12_-v2-h8/d8. **C-E**) GIRK current recordings of SNAP-mGluR2 photoswitching reveals reversibility and repeatability of switching, and increased performance of BGAG_12_-v2-d8 in terms of efficiency (D) and kinetics (E). P-values from unpaired t-tests reported in panels D and E.

In summary, we have translated the deuteration strategy from fluorophores to azobenzene photoswitches, where we find substantially improved properties. Future work is needed to fully decipher the underlying photophysical mechanism. We demonstrate the ability of deuteration to enhance azobenzene photoswitching on two distinct systems, a soluble photochromic ligand (AQ) and a tethered PORTL (BGAG), suggesting that this strategy can be widely applicable to the many azobenzene scaffolds and ligands which have been reported as photopharmacological tools. Interestingly, while all three compounds reported here showed clearly improved photoswitch kinetics, there was variability in the extent of effects on light absorbance and photoswitch efficiency, motivating future analyses of the vast chemical space for deuteration (or semi-deuteration) on the aromatic units of an azobenzene and/or their substituents (e.g. *N*-methyl amine deuteration) to further optimize this strategy.

## MATERIALS and METHODS

### Chemical synthesis

Chemical synthesis and characterization procedures are reported in the Supporting Information.

### Cell culture, molecular biology and patch clamp electrophysiology

HEK293 cells were cultured in Dulbecco’s Modified Eagle Medium (DMEM; Corning) supplemented with 10% fetal bovine serum (FBS) and maintained at 37º C and 5% CO_2_. Cells were seeded at low density in poly-L-lysine coated 18 mm coverslips and transfected the following day with Lipofectamine 2000 (Thermo Fisher Scientific). Plasmid expressing BK channel human alpha subunit (pBNJ13-hSlo)^42^ was kindly gifted by Prof. Teresa Giraldez (University of La Laguna, Spain). This construct was used for testing AQ compounds. For BGAG recordings, SNAP-mGluR2^43^, GIRK1-F137S^44^ and tdTomato as a transfection marker were co-transfected in cells in a 1:1:0.2 ratio.

Whole cell patch clamp recordings were performed 24 hr after transfection using an Axopatch 200B amplifier and a Digidata 1550B interface controlled by pClampex software (Molecular Devices). Recordings were performed in a bath solution containing (in mM): 120 KCl, 25 NaCl, 10 HEPES, 2 CaCl_2_, 1 MgCl_2_. Pipettes of 3-5 MΩ resistance were filled with intracellular solution (in mM: 140 KCl, 10 HEPES, 5 EGTA, 3 MgCl_2_, 3 Na_2_ATP, 0.2 Na_2_GTP). For AQ compounds, AQ-h8 and AQ-d8 were added to a final concentration of 1 mM in the pipette solution. For BGAG, cells were labelled with 1 μM of BGAG_12_-v2-h8 or BGAG_12_-v2-d8 for 45 min at 37ºC in extracellular solution. Labeling efficiency was measured using a fluorophore competition assay as previously.^45^ Photoactivation of the compounds was obtained through a computer controlled CoolLED pE-4000 attached to an inverted microscope and through a 40x objective. Light intensities at the focal plane were (in mW/mm^2^): 5.6 for 385 nm and 4.9 for 525 nm. For AQ compound photoswitching, a 0.1% neutral density ND filter (Chroma) was added to the 385 nm illumination path to produce lower light conditions for kinetics analysis (5.57 μW/mm^2^).

To obtain an I-V curve for BK channel activation, a step protocol of 50 ms of pulse ranging from -100 mV to + 200 mV in +20 mV increments was recorded. This was done in the presence of either wavelength (385 nm or 525 nm) throughout each sweep. Steady-state current at the end of the pulse, normalized to the maximum current observed in each individual cell, was plotted against the voltage applied to each step. The protocol for photoswitching of AQ compounds consisted of a voltage clamp of the cell at + 60 mV and applying pulses of 20 s of 385 nm immediately followed by 10 s of 525 nm light. For BGAG recordings, photoactivation by 385 nm was performed until the mGluR2 evoked GIRK current was in a steady state and after that, was quickly switched off by 525 nm light. Photoswitch efficiency was calculated as the amplitude of the 385 nm evoked current divided by the amplitude of the current response to saturating 1 mM glutamate.

All cellular data comes from at least three separate transfections/experimental days. Data was analyzed using Clampfit (Molecular Devices) and Prism 9 (GraphPad). AQ and BGAG trans-to-cis kinetics were quantified by fitting the evoked currents to a single exponential.

## ACKNOWLEDGEMENTS

This project has received funding from the European Union’s Horizon Europe Framework Programme (deuterON, grant agreement no. 101042046 to JB). A.G-H. is funded by the Margarita Salas Fellowship from the Spanish Ministry of Universities. J.L. is supported by the Rohr Family Research Scholar Award and the Irma T. Hirschl and Monique Weill-Caulier Award. We thank Aditi Jain for technical assistance.

