## Supporting Information for "Deuteration provides a general strategy to enhance azobenzene-based photopharmacology"

[1] Leibniz-Forschungsinstitut für Molekulare Pharmakologie (FMP), 13125 Berlin, Germany.

[2] Department of Biochemistry, Weill Cornell Medicine, New York, NY 10065, USA

[3] Department of Chemistry and Molecular Biology, University of Gothenburg, 413 90 Gothenburg, Sweden

[#] Equal contribution

* Correspondence should be addressed to:


**Table of Contents**

**1. General** ........ **3**

**2. SynthesisS** ........ **4**

*2.1. (E)-1,2-Diphenyldiazene (AB-h10)* *4*

*2.2. (E)-1,2-Bis(phenyl-$d_5$)diazene (AB-d10)* *4*

*2.3. (E)-4,4'-(Diazene-1,2-diyl)bis(benzen-2,3,5,6-$d_4$-amine) (2)* *5*

*2.4. (E)-2-((4-((4-Aminophenyl-2,3,5,6-$d_4$)diazenyl)phenyl-2,3,5,6-$d_4$)amino)-N,N,N-triethyl-2-oxoethan-1-aminium* *5*

*2.5. (E)-2-((4-((4-Acetamidophenyl-2,3,5,6-$d_4$)diazenyl)phenyl-2,3,5,6-$d_4$)amino)-N,N,N-triethyl-2-oxoethan-1-aminium (AQ-d8)* *6*

*2.6. (Dimethyl (2S,4S)-2-(4-((4-((E)-(4-aminophenyl-2,3,5,6-d4)diazenyl)phenyl-2,3,5,6-d4)amino)-4-oxobutyl)-4-((tert-butoxycarbonyl)amino)pentanedioate (4)* *6*

*2.7. Dimethyl (2S,4S)-2-(4-((4-((E)-(4-(2-aminoacetamido)phenyl-2,3,5,6-d4)diazenyl)phenyl-2,3,5,6-d4)amino)-4-oxobutyl)-4-((tert-butoxycarbonyl)amino)pentanedioate (5)* *7*

*2.8. Dimethyl (2S,4S)-2-(4-((4-((E)-(4-(1-amino-39-oxo-3,6,9,12,15,18,21,24,27,30,33,36-dodecaoxa-40-azadotetracontan-42-amido)phenyl-2,3,5,6-d4)diazenyl)phenyl-2,3,5,6-d4)amino)-4-oxobutyl)-4-((tert-butoxycarbonyl)amino)pentanedioate (6)* *8*

*2.9. (E)-2-((4-((4-Acetamidophenyl-2,3,5,6-d4)diazenyl)phenyl-2,3,5,6-d4)amino)-N,N,N-triethyl-2-oxoethan-1-aminium Dimethyl (2S,4S)-2-(4-((4-((E)-(4-(1-(4-(((2-amino-9H-purin-6-yl)oxy)methyl)phenyl)-3,7,47-trioxo-11,14,17,20,23,26,29,32,35,38,41,44-dodecaoxa-2,8,48-triazapentacontan-50-amido)phenyl-2,3,5,6-d4)diazenyl)phenyl-2,3,5,6-d4)amino)-4-oxobutyl)-4-((tert-butoxycarbonyl)amino)pentanedioate (7)* *8*

*2.10. (2S,4S)-2-Amino-4-(4-((4-((E)-(4-(1-(4-(((2-amino-9H-purin-6-yl)oxy)methyl)phenyl)-3,7,47-trioxo-11,14,17,20,23,26,29,32,35,38,41,44-dodecaoxa-2,8,48-triazapentacontan-50-amido)phenyl-2,3,5,6-d4)diazenyl)phenyl-2,3,5,6-d4)amino)-4-oxobutyl)pentanedioic acid (BGAG$_{12}$-v2-d8)* *9*

*2.11. (E)-2-((4-((4-Acetamidophenyl-2,3,5,6-d4)diazenyl)phenyl-2,3,5,6-d4)amino)-N,N,N-triethyl-2-oxoethan-1-aminium Dimethyl (2S,4S)-2-(4-((4-((E)-(4-(1-(4-(((2-amino-9H-purin-6-yl)oxy)methyl)phenyl)-3,7,47-trioxo-11,14,17,20,23,26,29,32,35,38,41,44-dodecaoxa-2,8,48-triazapentacontan-50-amido)phenyl)diazenyl)phenyl)amino)-4-oxobutyl)-4-((tert-butoxycarbonyl)amino)pentanedioate (8)* *10*

*2.12. (2S,4S)-2-Amino-4-(4-((4-((E)-(4-(1-(4-(((2-amino-9H-purin-6-yl)oxy)methyl)phenyl)-3,7,47-trioxo-11,14,17,20,23,26,29,32,35,38,41,44-dodecaoxa-2,8,48-triazapentacontan-50-amido)phenyl)diazenyl)phenyl)amino)-4-oxobutyl)pentanedioic acid (BGAG$_{12}$-v2-h8)* *11*

**3. NMR spectroscopy** ........ **12**

*3.1. (E)-1,2-Bis(phenyl-d5)diazene* *12*

*3.2. (E)-1,2-Diphenyldiazene* *13*

*3.3. (E)-4,4'-(Diazene-1,2-diyl)bis(benzen-2,3,5,6-$d_4$-amine)* *14*

*3.4. (E)-2-((4-((4-Aminophenyl-2,3,5,6-$d_4$)diazenyl)phenyl-2,3,5,6-$d_4$)amino)-N,N,N-triethyl-2-oxoethan-1-aminium* *15*

*3.5. (E)-2-((4-((4-Acetamidophenyl-2,3,5,6-$d_4$)diazenyl)phenyl-2,3,5,6-$d_4$)amino)-N,N,N-triethyl-2-oxoethan-1-aminium* *16*

### 1. General

All chemical reagents and anhydrous solvents for synthesis were purchased from commercial suppliers (Sigma-Aldrich, Fluka, Acros, Fluorochem, TCI) and were used without further purification.

NMR spectra were recorded in deuterated solvents on a Bruker AVANCE III 600 equipped with a CryoProbe or on a Bruker AVANCE II 750 and calibrated to residual solvent peaks ($^{1}H/^{13}C$ in ppm): DMSO-$d_6$ (2.50/39.52), MeOD-$d_4$ (3.31/49.00). Multiplicities are abbreviated as follows: s = singlet, d = doublet, t = triplet, q = quartet, p = pentet, h = heptet, br = broad, m = multiplet. Coupling constants *J* are reported in Hz. Spectra are reported based on appearance, not on theoretical multiplicities derived from structural information. Light for switching was delivered by a CoolLED pE-4000 (intensities used: $I_{385\ nm}$ = 0.459 mW/mm$^2$; $I_{500\ nm}$ = 8.20 mW/mm$^2$; $I_{525\ nm}$ = 4.95 mW/mm$^2$). Full spectra were acquired either in the dark or under appropriate illumination, and photostationary states were recorded at one wavelength under annotated illumination.

UPLC-UV/Vis for purity assessment was performed on an Agilent 1260 Infinity II LC System equipped with Agilent SB- C18 column (1.8 µm, 2.1 × 50 mm). Buffer A: 0.1% FA in H2O Buffer B: 0.1% FA acetonitrile. The typical gradient was from 10% B for 0.5 min → gradient to 95% B over 5 min → 95% B for 0.5 min → gradient to 99% B over 1 min with 0.8 mL/min flow. Retention times (tR) are given in minutes (min). Chromatograms were imported into Graphpad Prism10 and plotted.

High resolution ESI-MS spectra were recorded on a Waters H-class instrument equipped with a quaternary solvent manager, a Waters sample manager-FTN, a Waters PDA detector and a Waters column manager with an Acquity UPLC protein BEH C18 column (1.7 µm, 2.1 mm x 50 mm). Samples were eluted with a flow rate of 0.3 mL/min. The following gradient was used: A: 0.01 % FA in H2O; B: 0.01 % FA in MeCN. 5 % B: 0-1 min; 5 to 95 % B: 1-7min; 95 % B: 7 to 8.5 min. Mass analysis was conducted with a Waters XEVO G2-XS QTof analyzer.

Reactions and chromatography fractions were monitored by thin layer chromatography (TLC) on Merck silica gel 60 F254 glass plates. The spots were visualized either under UV light at 254 nm and/or 366 nm or with appropriate staining method (iodine, *para*-anisaldehyde, $KMnO_4$) followed by heating.

UV/Vis spectroscopy was performed on either: i) a JASCO V-550 UV/Vis/NIR spectrophotometer or ii) a Thermo ScientificTM NanoDropTM OneC microvolume-UV/Vis-spectrophotometer using a Hellma quartz glass cuvette (10 mm pathlength). Light for switching was delivered by a CoolLED pE-4000 (intensities used: $I_{385\ nm}$ = 0.459 mW/mm$^2$; $I_{500\ nm}$ = 8.20 mW/mm$^2$; $I_{525\ nm}$ = 4.95 mW/mm$^2$). Full spectra were acquired either in the dark or under appropriate illumination, and kinetic traces were recorded at 365 nm under annotated illumination. Results were exported and plotted in GraphPad Prism 10 and kinetic traces were fitted monoexponentially GraphPad Prism 10 and plotted in GraphPad Prism 10.

IR spectroscopy was performed on a Bruker Tensor 27 FT-IR and evaluated with OPUS 8.5.

### 2. SynthesisS

#### 2.1. (*E*)-1,2-Diphenyldiazene (AB-h10)

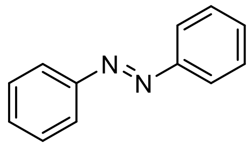

A 50 mL flask was charged with nitrobenzene (96 mg, 781 µmol, 1.0 equiv.) in 10 mL methanol, and a solution of sodium hydroxide (101.5 mg, 1.56 mmol, 2.0 equiv.) dissolved in 3 mL $H_2O$ was added, before zinc dust (125.0 mg, 3.12 mmol, 4.0 equiv.) was added and the reaction was stirred for 12 h under reflux. The mixture was filtered while hot, and concentrated *in vacuo*. The residue was again dissolved in 2 mL hot methanol, and dried onto isolute. The mixture was subjected to column chromatography, with a mixture of EtOAc/cyclohexane (1:99). The product was concentrated *in vacuo* to obtain 37.0 mg (203 µmol) of the desired compound as an orange powder in 52% yield.

**$^1$H NMR** (600 MHz, MeOD): $\delta$ [ppm] = 7.93 (d, $J$ = 7.4 Hz, 4H), 7.57 (t, $J$ = 7.4 Hz, 4H), 7.53 (t, $J$ = 7.2 Hz, 2H).
**$^{13}$C NMR** (150 MHz, MeOD): $\delta$ [ppm] = 152.6, 130.9, 128.9, 122.4.
**HRMS** (ESI): calc. for $C_{12}H_{10}N_2$ $[M+H]^+$: 183.0917, found: 183.0956.
**IR** (wavenumber / cm$^{-1}$): 3063.7, 2925.3, 1957.4, 1899.0, 1808.2, 1771.3, 1685.4, 1582.2, 1539.7, 1483.1, 1453.3, 1398.0, 1299.4, 1221.7, 1151.2, 1071.2, 1019.7, 999.7, 985.4, 926.0, 851.2

#### 2.2. (*E*)-1,2-Bis(phenyl-$d_5$)diazene (AB-d10)

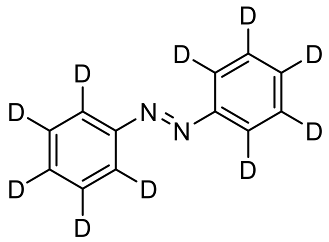

A 50 mL flask was charged with 1-nitrobenzene-2,3,4,5,6-$d_5$ (100 mg, 781 µmol, 1.0 equiv.) in 10 mL methanol, and a solution of sodium hydroxide (101.5 mg, 1.56 mmol, 2.0 equiv.) dissolved in 3 mL $H_2O$ was added, before zinc dust (125.0 mg, 3.12 mmol, 4.0 equiv.) was added and the reaction was stirred for 12 h under reflux. The mixture is filtered while hot, and concentrated *in vacuo*. The residue was again dissolved in 2 mL hot methanol, and put on isolute. The mixture was subjected to column chromatography, with a mixture of EtOAc/cyclohexane (1:99). The product was concentrated *in vacuo* to obtain 34.0 mg (177 µmol) of the desired compound as an orange powder in 45% yield.

**$^1$H NMR** (600 MHz, MeOD): $\delta$ [ppm] = no peaks observed.
**$^{13}$C NMR** (150 MHz, MeOD): $\delta$ [ppm] = 152.5, 130.5, 128.5, 122.0.
**HRMS** (ESI): calc. for $C_{12}D_{10}N_2$ [M+H]+: 193.1544, found: 193.1531
**IR** (wavenumber / cm$^{-1}$): 3853.9, 3838.6, 3735.6, 3670.2, 3648.3, 2926.3, 2853.3, 2270.5, 1733.6, 1616.6, 1541.6, 1457.5, 1364.3, 1294.1, 1168.7, 1080.0, 1035.7, 958.8, 868.8, 823.4, 811.3

### 2.3. (*E*)-4,4'-(Diazene-1,2-diyl)bis(benzen-2,3,5,6-d$_4$-amine) (2)

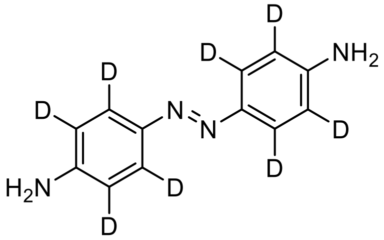

A round bottom flask was charged with benzene-d$_4$-1,4-diamine (**1**, 100 mg, 893 µmol, 1.0 equiv.) and dissolved in 50 mL $CH_2Cl_2$. To the vigorously stirred solution was added Dess-Martin-Periodinane (832 mg, 1.96 mmol, 2.2 equiv.) dissolved in 50 mL $CH_2Cl_2$ over 90 min, and the mixture was stirred for an additional 30 min. The mixture was subjected directly to column chromatography to first elute $CH_2Cl_2$, before the residue was purified with a mixture of EtOAc/cyclohexane (50:50). The solution was concentrated *in vacuo* to obtain 28.0 mg (127 µmol) of the desired compound as an orange powder in 28% yield.

**$^1$H NMR** (600 MHz, DMSO-d$_6$): $\delta$ [ppm] = no peaks observed.
**$^2$H NMR** (92 MHz, DMSO-d$_6$): $\delta$ [ppm] = 7.54 (s, 4H), 6.64 (s, 4H).
**$^{13}$C NMR** (150 MHz, DMSO-d$_6$): $\delta$ [ppm] = 148.2, 145.0, 124.0, 115.6.
**HRMS** (ESI): calc. for $C_{12}H_4D_8N_4$ $[M+H]^+$: 221.1637, found: 221.1647.

### 2.4. (*E*)-2-((4-((4-Aminophenyl-2,3,5,6-d$_4$)diazenyl)phenyl-2,3,5,6-d$_4$)amino)-*N,N,N*-triethyl-2-oxoethan-1-aminium

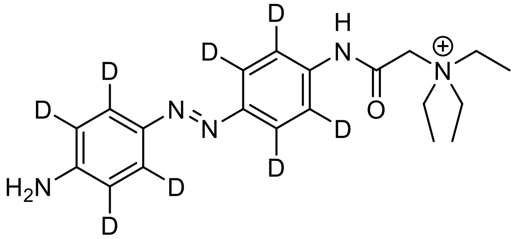

A 50 mL round bottom flask was charged with *E*)-4,4'-(diazene-1,2-diyl)bis(benzen-2,3,5,6-d$_4$-amine (**2**, 14 mg, 63.6 µmol, 1.0 equiv.), *N*-(carboxymethyl)-*N,N*-diethylethanaminium chloride (Banghart et al., 2009) (30.5 mg, 191 µmol, 3.0 equiv.), dissolved in 1 mL DMSO, before 86 µL DIPEA was added. To the vigorously stirred solution was added HBTU (79.5 mg 210 µmol, 3.3 equiv.) in one portion and the reaction mixture was stirred at 50 °C o.n. before it was quenched by addition of 200 µL HOAc and 200 µL $H_2O$ and subjected to RP-HPLC purification to obtain 5.9 mg (16.3 µmol) of the desired product in 26% yield, while 43% of the starting material were recovered.

**$^1$H NMR** (600 MHz, DMSO-d$_6$): $\delta$ [ppm] = 10.9 (s, 1H), 4.20 (s, 2H), 3.55 (q, $J$ = 7.2 Hz, 6H), 1.28 (t, $J$ = 7.2 Hz, 9H).
**$^{13}$C NMR** (150 MHz, DMSO-d$_6$): $\delta$ [ppm] = 162.4, 152.8, 149.3, 143.2, 138.9, 124.9, 122.4, 120.1, 113.6, 56.8, 54.5, 7.88.
**HRMS** (ESI): calc. for $C_{20}H_{20}D_8N_5O^+$ $[M]^+$: 362.2791, found: 362.2758.

#### 2.5. (*E*)-2-((4-((4-Acetamidophenyl-2,3,5,6-d$_4$)diazenyl)phenyl-2,3,5,6-d$_4$)amino)-*N,N,N*-triethyl-2-oxoethan-1-aminium (AQ-d8)

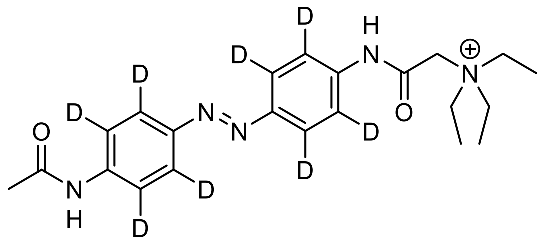

A flame-dried 50 mL Schlenk flask under an argon atmosphere was charged with (*E*)-2-((4-((4-aminophenyl-2,3,5,6-d$_4$)diazenyl)phenyl-2,3,5,6-d$_4$)amino)-*N,N,N*-triethyl-2-oxoethan-1-aminium (1.5 mg, 4.14 µmol, 1.0 equiv.) and dissolved in 1 mL dry THF, before 1.31 µL DIPEA (8.28 µmol, 2.0 equiv.) was added. The reaction mixture was vigorously stirred and cooled to 0 °C in an ice-bath, before AcCl (0.5 µL, 6.21 µmol, 1.5 equiv.) was added. Upon addition, the ice bath was removed and the reaction mixture was allowed to stir for 30 min while warming to r.t., before it was quenched by addition of glacial HOAc (10 µL). The reaction mixture was concentrated *in vacuo* and subjected to RP-HPLC to obtain 0.9 mg (2.23 µmol) of the desired compound (TFA salt) as a highly hygroscopic, orange powder in 54% yield.

**$^1$H NMR** (600 MHz, $D_2O$): $\delta$ [ppm] = 4.09 (s, 2H), 3.57 (q, $J$ = 7.2 Hz, 6H), 1.32 (t, $J$ = 7.2 Hz, 9H).
**$^{13}$C NMR** (150 MHz, $D_2O$): $\delta$ [ppm] = 56.5, 54.7, 23.1, 7.0.
**HRMS** (ESI): calc. for $C_{22}H_{22}D_8N_5O_2^+$ $[M]^+$: 404.2897, found: 404.2898.
**IR** (wavenumber / cm$^{-1}$): 3838.4, 3750.4, 3648.9, 2365.3, 1670.1, 1567.9, 1521.5, 1456.9, 1417.9, 1372.8, 1318.0, 1245.3, 1200.1, 1127.4, 1007.3, 833.4

**IR** (wavenumber / cm$^{-1}$) for AQ-h8 (ref[1]): 3936.1, 3862.8, 3838.4, 3802.6, 3751.5, 3734.1, 3670.9, 3332.7, 2364.6, 1684.5, 1541.1, 1115.9, 1078.9, 1012.1, 842.5

#### 2.6. (Dimethyl (2*S*,4*S*)-2-(4-((4-((*E*)-(4-aminophenyl-2,3,5,6-d4)diazenyl)phenyl-2,3,5,6-d4)amino)-4-oxobutyl)-4-((*tert*-butoxycarbonyl)amino)pentanedioate (4)

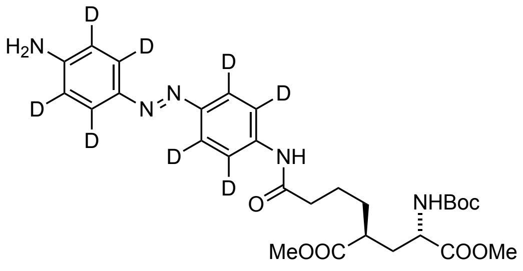

A round bottom flask was charged with (5*S*,7*S*)-7-((*tert*-butoxycarbonyl)amino)-8-methoxy-5-(methoxycarbonyl)-8-oxooctanoic acid (3, 32 mg, 90.9 µmol, 1.0 equiv.), 2 (20 mg, 90.9 µmol, 1.0 equiv.) and dissolved in DMSO (2 mL) and DIPEA (47 mg, 364 µmol, 63 µL, 4.0 equiv.) before HBTU (35 mg, 90.9 µmol, 1.0 equiv.) was added. The reaction mixture was allowed to incubate at r.t. for 4 hours before it was quenched with HOAc (63 µL) and water (100 µL) and subjected to RP-HPLC purification. The product containing fractions were pooled and lyophilized to obtain 21 mg (37.3 µmol) of the desired product in 41% yield as a red powder.

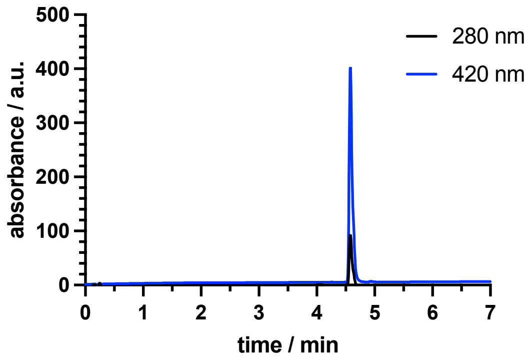

**LRMS** (ESI): calc. for $C_{28}H_{30}D_8N_5O_7^+$ $[M+H]^+$: 564.3, found:564.3.

#### 2.7. Dimethyl (2*S*,4*S*)-2-(4-((4-((*E*)-(4-(2-aminoacetamido)phenyl-2,3,5,6-d4)diazenyl)phenyl-2,3,5,6-d4)amino)-4-oxobutyl)-4-((*tert*-butoxycarbonyl)amino)pentanedioate (5)

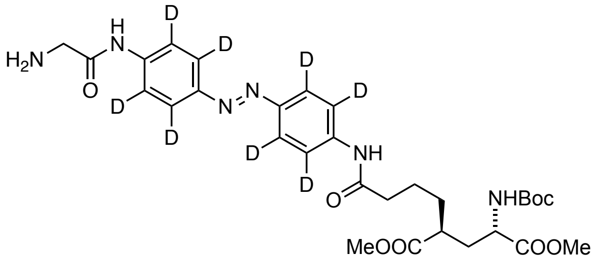

A round bottom flask was charged with (4, 21 mg, 37 µmol, 1.0 equiv.), dissolved in dry THF (7 mL) and DIPEA (18 mg, 142 µmol, 25 µL, 4.0 equiv.) and was cooled to 0 °C before FmocGlyCl (22.4 mg, 71 µmol, 1.9 equiv.) was added. The reaction mixture was allowed to incubate at r.t. for 1 hour before DBU in MeCN (150 µL in 5 mL) was added. After 3 hours, the reaction was quenched with HOAc (200 µL) and water (1 mL) and subjected to RP-HPLC purification. The product containing fractions were pooled and lyophilized to obtain 10 µmol of the desired product in 27% yield as an orange powder.

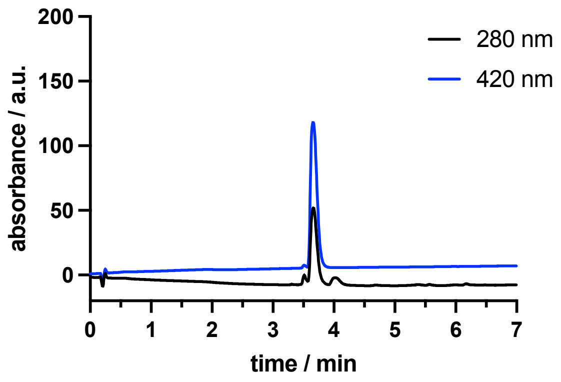

**LRMS** (ESI): calc. for $C_{30}H_{33}D_8N_6O_8^+$ $[M+H]^+$: 621.3, found: 621.4.

#### 2.8. Dimethyl (2*S*,4*S*)-2-(4-((4-((*E*)-(4-(1-amino-39-oxo-3,6,9,12,15,18,21,24,27,30,33,36-dodecaoxa-40-azadotetracontan-42-amido)phenyl-2,3,5,6-d4)diazenyl)phenyl-2,3,5,6-d4)amino)-4-oxobutyl)-4-((*tert*-butoxycarbonyl)amino)pentanedioate (6)

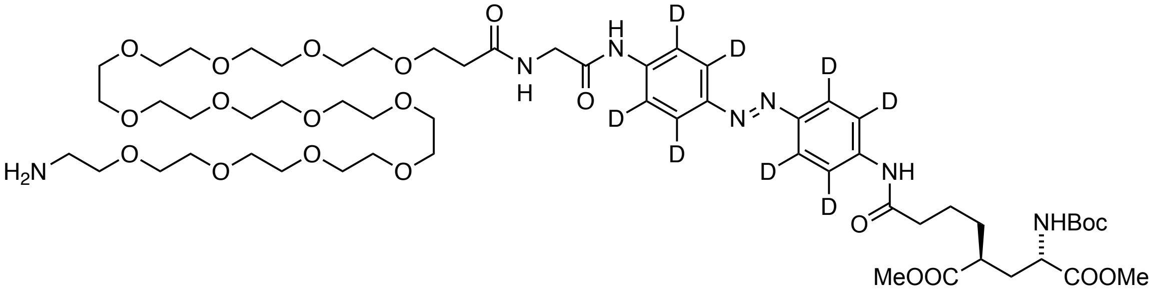

A 1.5 mL Eppendorf vial was charged with 5 (10 µmol, 1.0 equiv.), Fmoc-PEG$_{12}$-COOH (8.4 mg, 10 µmol, 1.0 equiv.) and dissolved in DMF (1 mL) and DIPEA (6.7 mg, 53 µmol, 9.3 µL, 5.3 equiv.) before HBTU (3.8 mg, 10 µmol, 1.0 equiv.) was added. The reaction mixture was allowed to incubate at r.t. for 5 hours before the addition of DBU (50 µL) and let stand for another hour. The reaction was quenched with HOAc (100 µL) and water (200 µL) and subjected to RP-HPLC purification. The product containing fractions were pooled and lyophilized to obtain 8.4 µmol of the desired product in 84% yield as an orange powder.

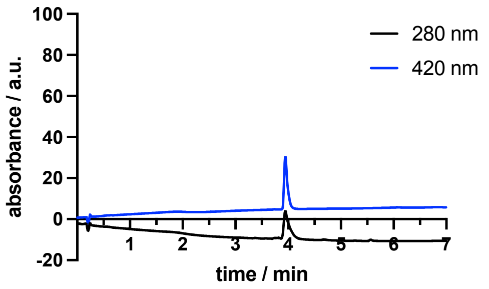

**LRMS** (ESI): calc. for $C_{57}H_{86}D_8N_7NaO_{21}^+$ $[M+H+Na]^{2+}$: 621.8, found: 622.0.

#### 2.9. (*E*)-2-((4-((4-Acetamidophenyl-2,3,5,6-d4)diazenyl)phenyl-2,3,5,6-d4)amino)-*N*,*N*,*N*-triethyl-2-oxoethan-1-aminium Dimethyl (2*S*,4*S*)-2-(4-((4-((*E*)-(4-(1-(4-(((2-amino-9*H*-purin-6-yl)oxy)methyl)phenyl)-3,7,47-trioxo-11,14,17,20,23,26,29,32,35,38,41,44-dodecaoxa-2,8,48-triazapentacontan-50-amido)phenyl-2,3,5,6-d4)diazenyl)phenyl-2,3,5,6-d4)amino)-4-oxobutyl)-4-((*tert*-butoxycarbonyl)amino)pentanedioate (7)

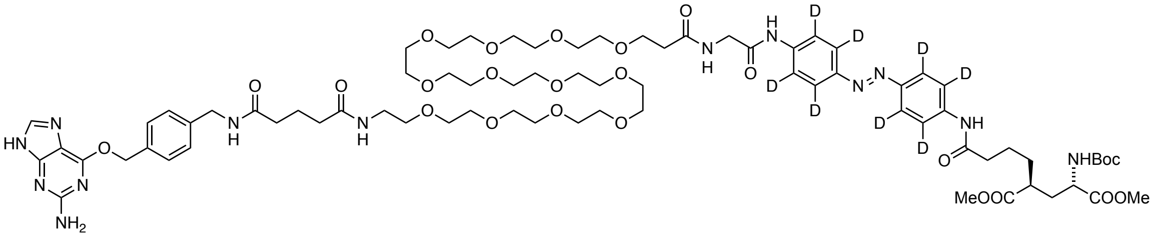

A 1.5 mL Eppendorf vial was charged with 6 (8.4 µmol, 1.0 equiv.), BG-COOH (4.5 mg, 12.6 µmol, 1.5 equiv.) and dissolved in DMF (1 mL) and DIPEA (8.7 mg, 67 µmol, 9.5 µL, 8.0 equiv.) before HBTU (4.8 mg, 12.6 µmol, 1.5 equiv.) was added. The reaction mixture was allowed to incubate at r.t. for 4 hours before it was quenched with HOAc (100 µL) and water (200 µL) and subjected to RP-HPLC purification. The product containing fractions were pooled and lyophilized to obtain 6.0 µmol of the desired product in 71% yield as an orange powder.

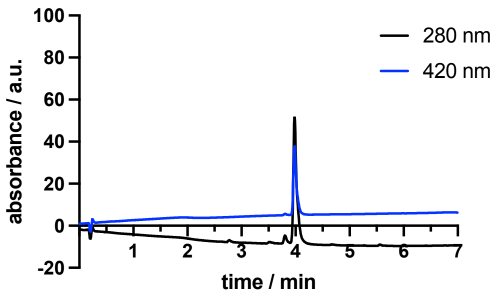

**LRMS** (ESI): calc. for $C_{75}H_{104}D_8N_{13}O_{24}^+$ [M+H]$^+$: 1586.8, found: 1586.6.

### 2.10. (2*S*,4*S*)-2-Amino-4-(4-((4-((*E*)-(4-(1-(4-(((2-amino-9*H*-purin-6-yl)oxy)methyl)phenyl)-3,7,47-trioxo-11,14,17,20,23,26,29,32,35,38,41,44-dodecaoxa-2,8,48-triazapentacontan-50-amido)phenyl-2,3,5,6-d4)diazenyl)phenyl-2,3,5,6-d4)amino)-4-oxobutyl)pentanedioic acid (BGAG$_{12}$-v2-d8)

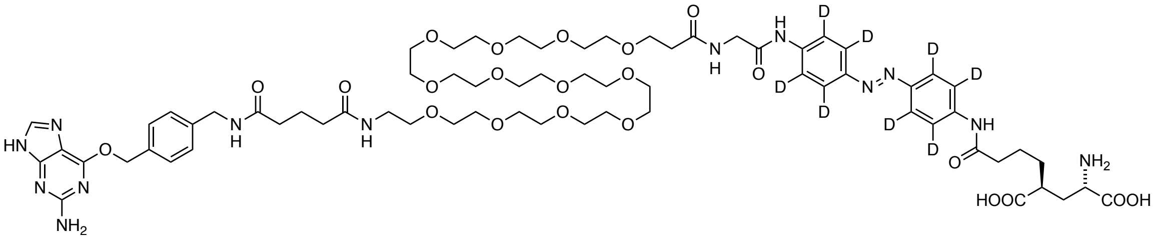

A 15 mL Falcon tube was charged with 7 (6.0 µmol, 1.0 equiv.) and MeOH (250 µL), THF (500 µL) and 1 M LiOH (250 µL) were added sequentially. The reaction mixture was allowed to incubate at r.t. for 2 hours, before volatiles were removed first by a gentle stream of nitrogen and the by lyophilization. The mixture was then cooled to 0 °C and ice-cold TFA (1 mL) was added and the reaction mixture turned deep purple and was allowed to incubate for 1 hour. Removal of the volatiles with a gentle stream of nitrogen by maintaining cooling to 0 °C was performed, before the residue was taken up in 1 mL of MeCN/$H_2O$/HOAc (25/25/1) and subjected to RP-HPLC purification. The product containing fractions were pooled and lyophilized to obtain 4.7 µmol of the desired product in 78% yield as an orange powder.

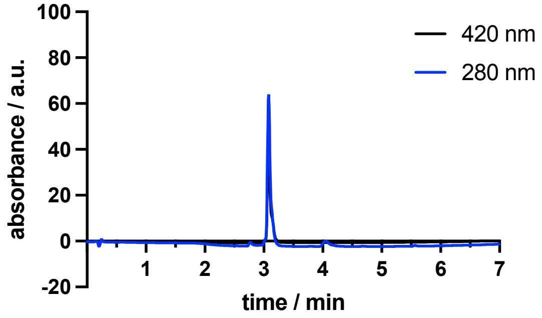

**LRMS** (ESI): calc. for $C_{68}H_{92}D_8N_{13}O_{22}^+$ [M+H]$^+$: 1458.8, found: 1459.6.

**2.11. (*E*)-2-((4-((4-Acetamidophenyl-2,3,5,6-d4)diazenyl)phenyl-2,3,5,6-d4)amino)-*N*,*N*,*N*-triethyl-2-oxoethan-1-aminium Dimethyl (2*S*,4*S*)-2-(4-((4-((*E*)-(4-(1-(4-(((2-amino-9*H*-purin-6-yl)oxy)methyl)phenyl)-3,7,47-trioxo-11,14,17,20,23,26,29,32,35,38,41,44-dodecaoxa-2,8,48-triazapentacontan-50-amido)phenyl)diazenyl)phenyl)amino)-4-oxobutyl)-4-((*tert*-butoxycarbonyl)amino)pentanedioate (8)**

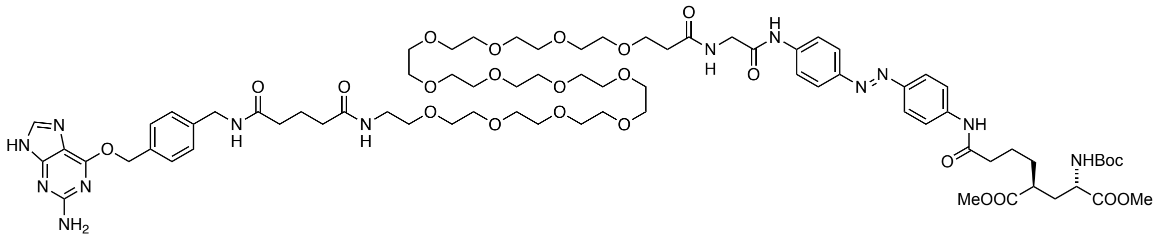

A 1.5 mL Eppendorf vial was charged with dimethyl (2*S*,4*S*)-2-(4-((4-((*E*)-(4-(1-amino-39-oxo-3,6,9,12,15,18,21,24,27,30,33,36-dodecaoxa-40-azadotetracontan-42-amido)phenyl)diazenyl)-phenyl)amino)-4-oxobutyl)-4-((*tert*-butoxycarbonyl)amino)pentanedioate (ref[2]) (22 µmol, 1.0 equiv.), BG-COOH (11.7 mg, 33 µmol, 1.5 equiv.) and dissolved in DMF (2 mL) and DIPEA (23 mg, 176 µmol, 31 µL, 8.0 equiv.) before HBTU (12.5 mg, 33 µmol, 1.5 equiv.) was added. The reaction mixture was allowed to incubate at r.t. for 4 hours before it was quenched with HOAc (100 µL) and water (300 µL) and subjected to RP-HPLC purification. The product containing fractions were pooled and lyophilized to obtain 12.2 µmol of the desired product in 55% yield as an orange powder.

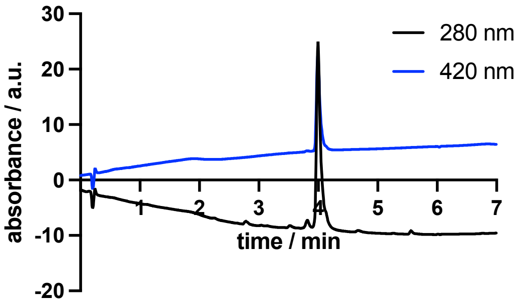

**LRMS** (ESI): calc. for $C_{75}H_{112}N_{13}O_{24}^{+}$ $[M+H]^{+}$: 1578.8, found: 1578.6.

**2.12. (2*S*,4*S*)-2-Amino-4-(4-((4-((*E*)-(4-(1-(4-(((2-amino-9*H*-purin-6-yl)oxy)methyl)phenyl)-3,7,47-trioxo-11,14,17,20,23,26,29,32,35,38,41,44-dodecaoxa-2,8,48-triazapentacontan-50-amido)phenyl)diazenyl)phenyl)amino)-4-oxobutyl)pentanedioic acid (BGAG$_{12}$-v2-h8)**

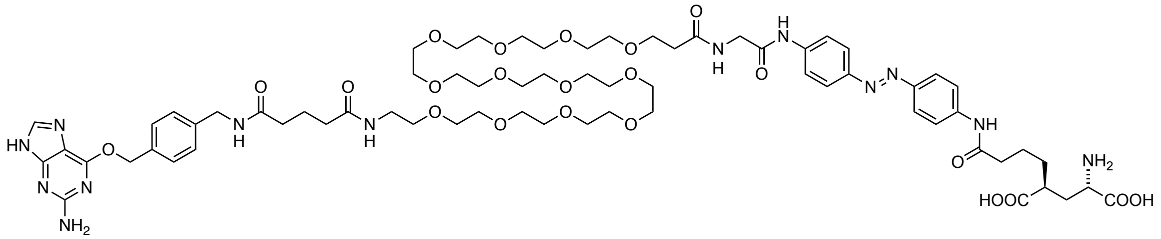

A 15 mL Falcon tube was charged with 8 (12.2 µmol, 1.0 equiv.) and MeOH (250 µL), THF (500 µL) and 1 M LiOH (250 µL) were added sequentially. The reaction mixture was allowed to incubate at r.t. for 2 hours, before volatiles were removed first by a gentle stream of nitrogen and the by lyophilization. The mixture was then cooled to 0 °C and ice-cold TFA (1 mL) was added and the reaction mixture turned deep purple and was allowed to incubate for 1 hour. Removal of the volatiles with a gentle stream of nitrogen by maintaining cooling to 0 °C was performed, before the residue was taken up in 1 mL of MeCN/$H_2O$/HOAc (25/25/1) and subjected to RP-HPLC purification. The product containing fractions were pooled and lyophilized to obtain 4.1 µmol of the desired product in 34% yield as an orange powder.

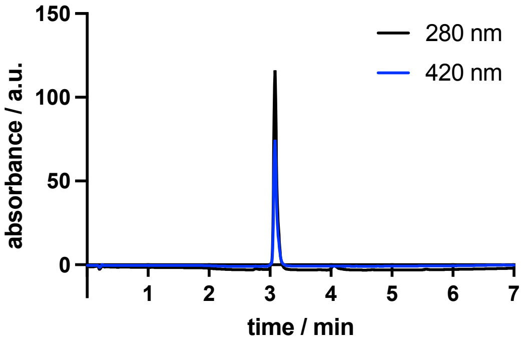

**LRMS** (ESI): calc. for $C_{68}H_{100}N_{13}O_{22}^{+}$ $[M+H]^{+}$: 1450.7, found: 1450.0.

### 3. NMR spectroscopy

#### 3.1. (*E*)-1,2-Bis(phenyl-d5)diazene

**$^1$H-NMR**

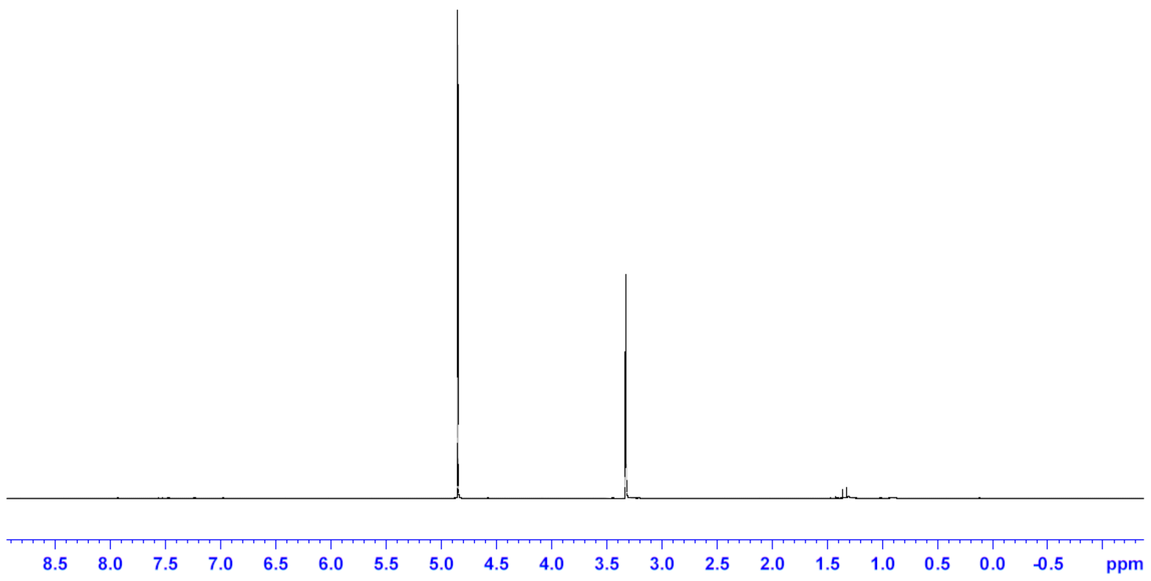

**$^{13}$C-NMR**

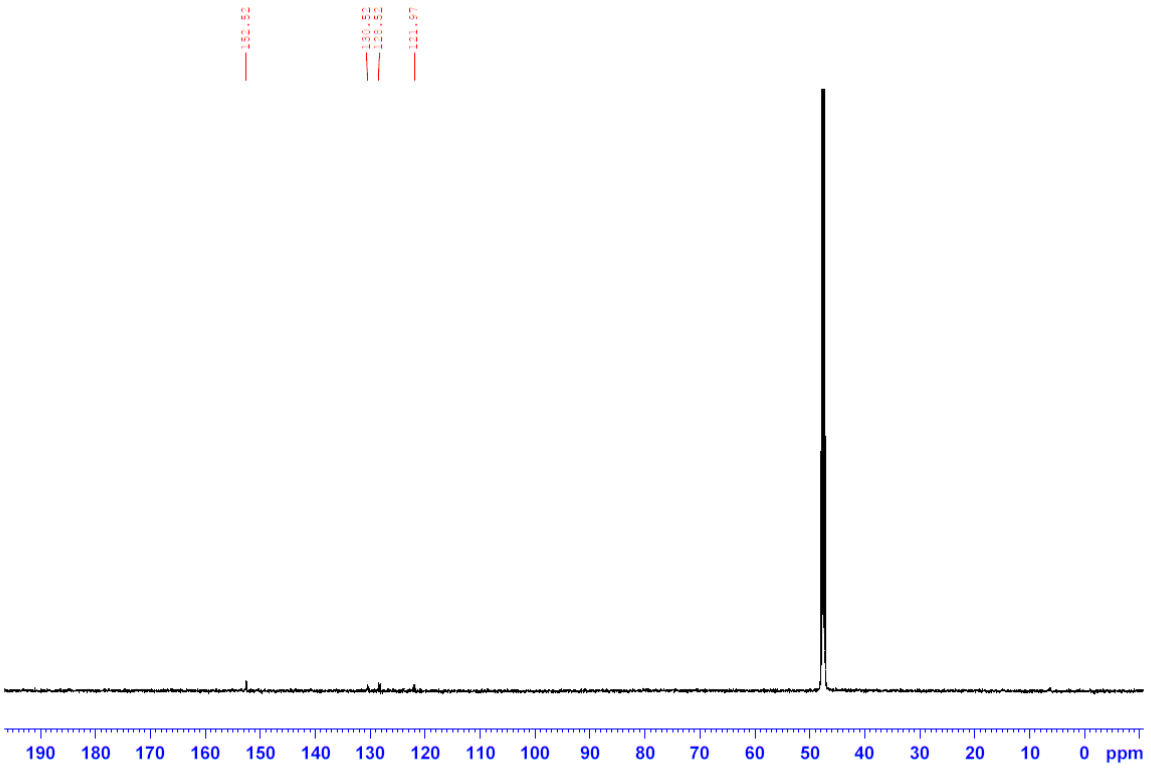

**3.2. (*E*)-1,2-Diphenyldiazene**

**[1]H-NMR**

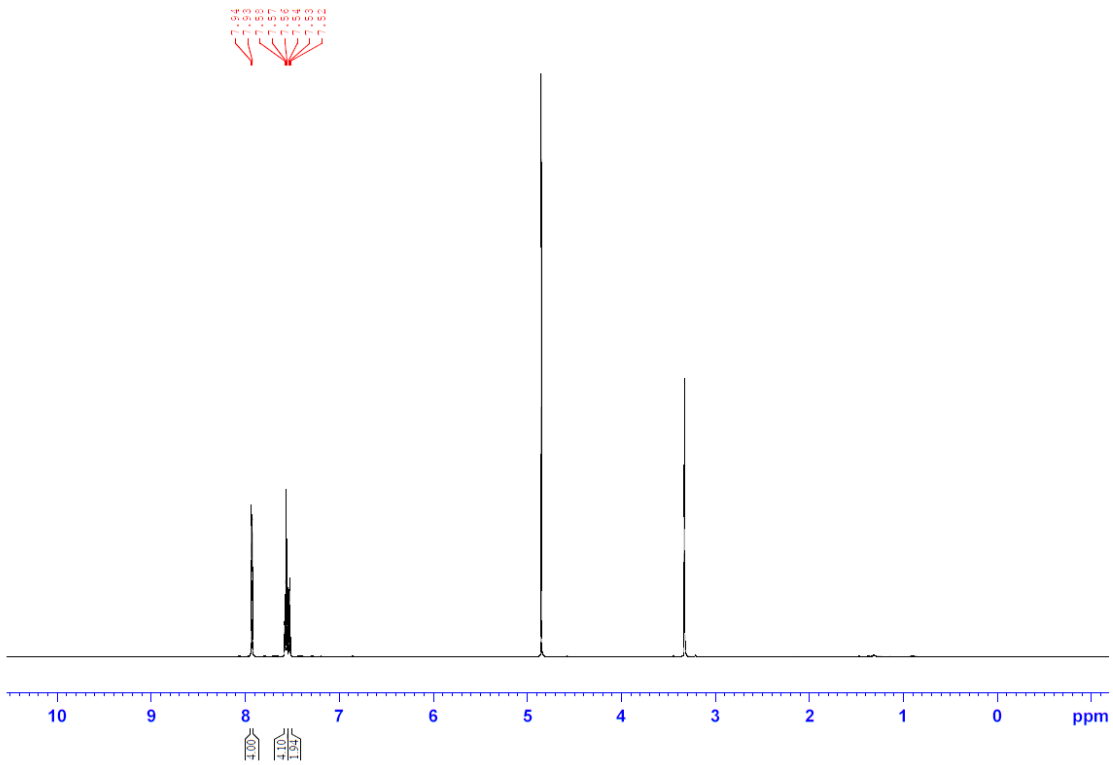

**[13]C-NMR**

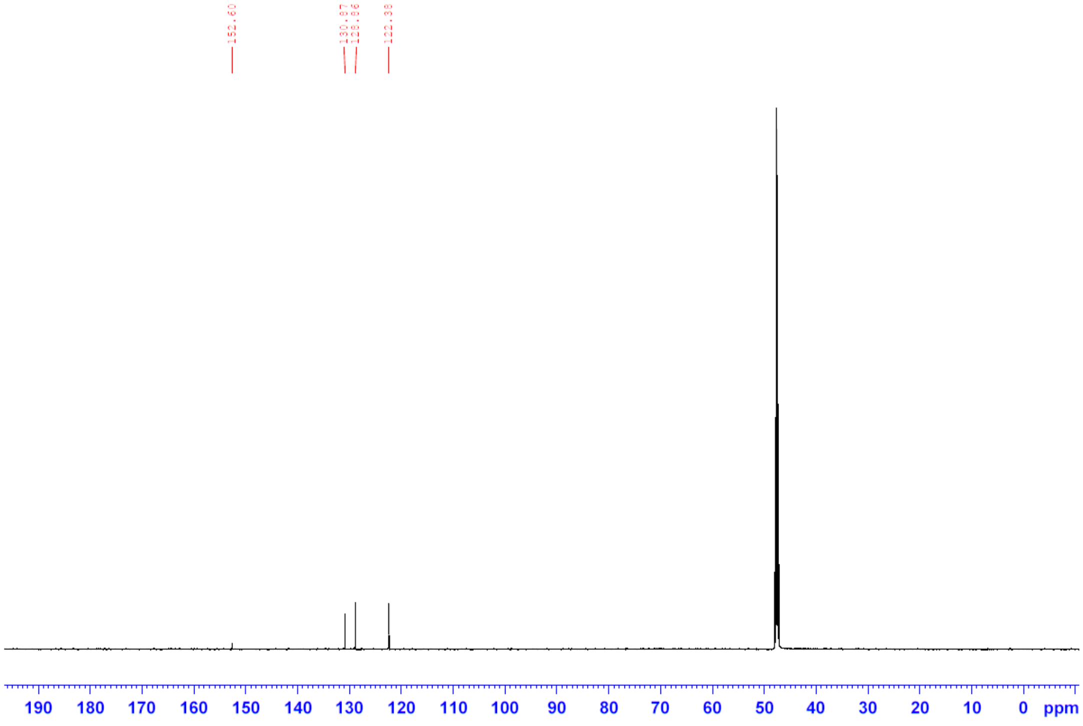

#### 3.3. (*E*)-4,4'-(Diazene-1,2-diyl)bis(benzen-2,3,5,6-d$_4$-amine)

**$^2$H-NMR**

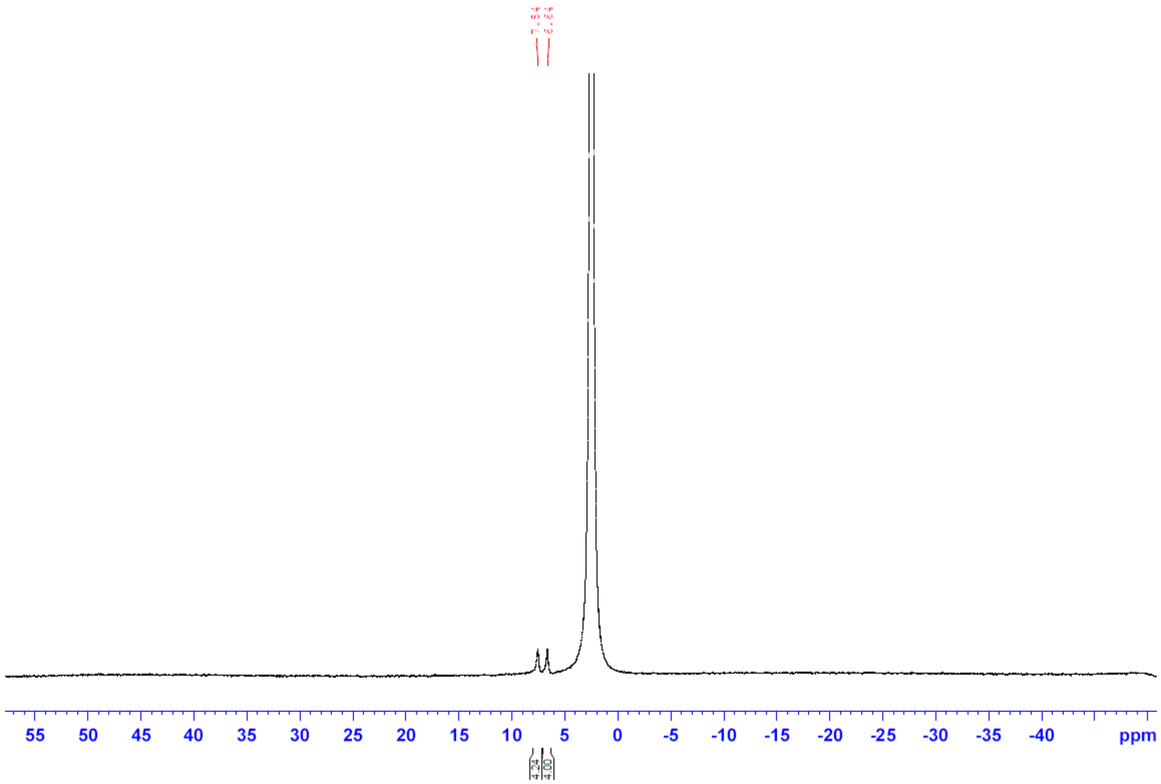

**$^{13}$C-NMR**

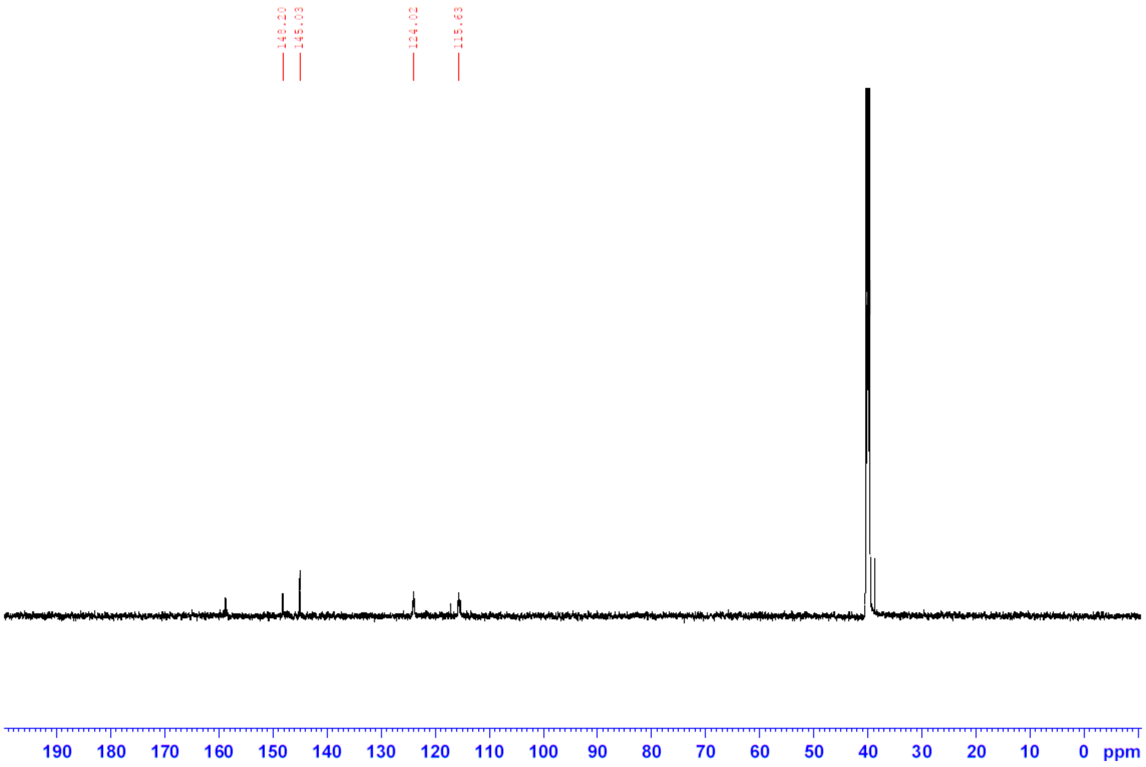

#### 3.4. (*E*)-2-((4-((4-Aminophenyl-2,3,5,6-d$_4$)diazenyl)phenyl-2,3,5,6-d$_4$)amino)-*N,N,N*-triethyl-2-oxoethan-1-aminium

**$^1$H-NMR**

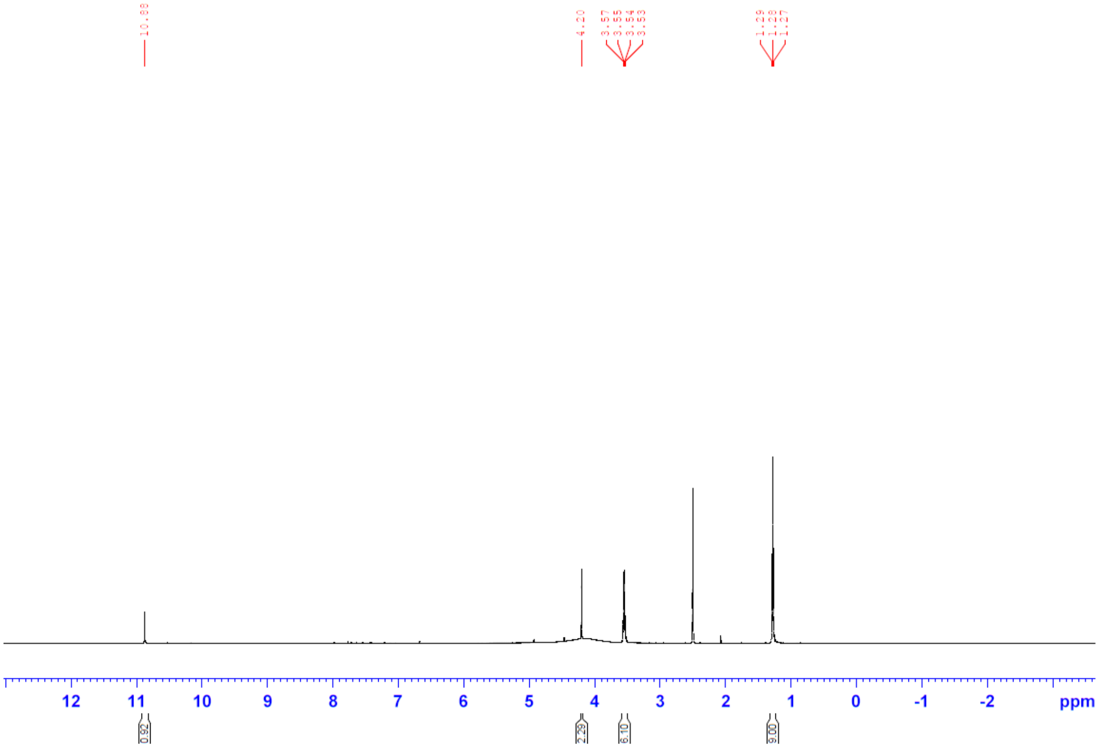

**$^{13}$C-NMR**

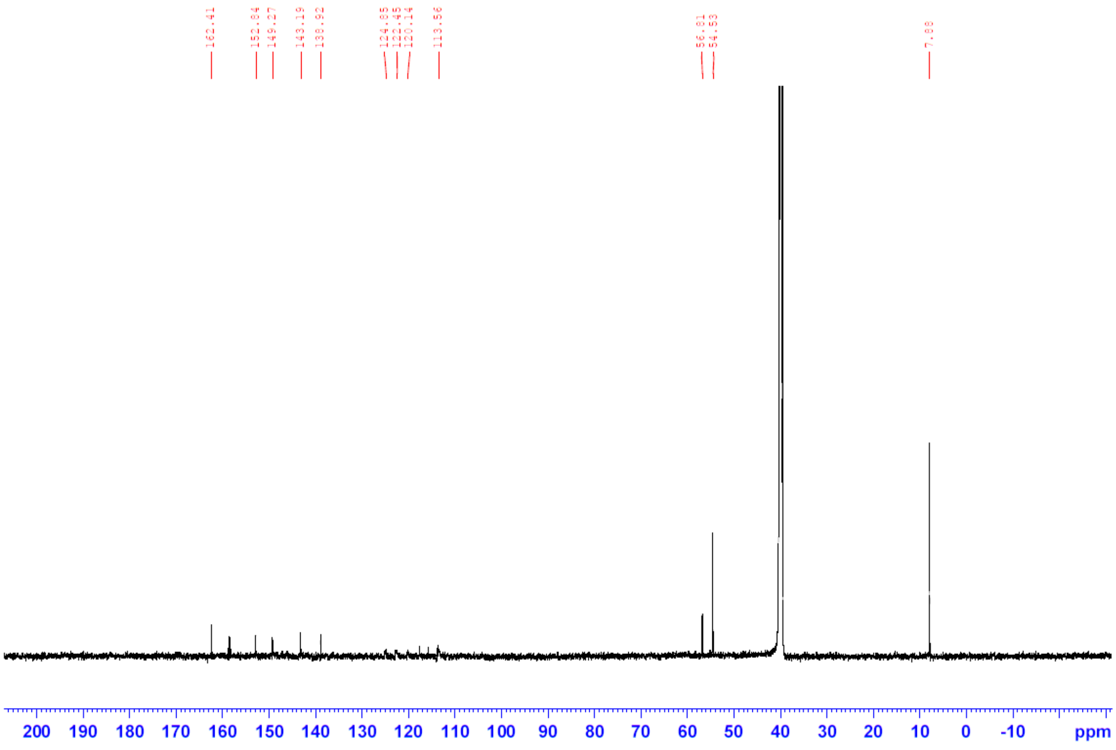

**3.5. (*E*)-2-((4-((4-Acetamidophenyl-2,3,5,6-d$_4$)diazenyl)phenyl-2,3,5,6-d$_4$)amino)-*N*,*N*,*N*-triethyl-2-oxoethan-1-aminium**

**$^1$H-NMR**

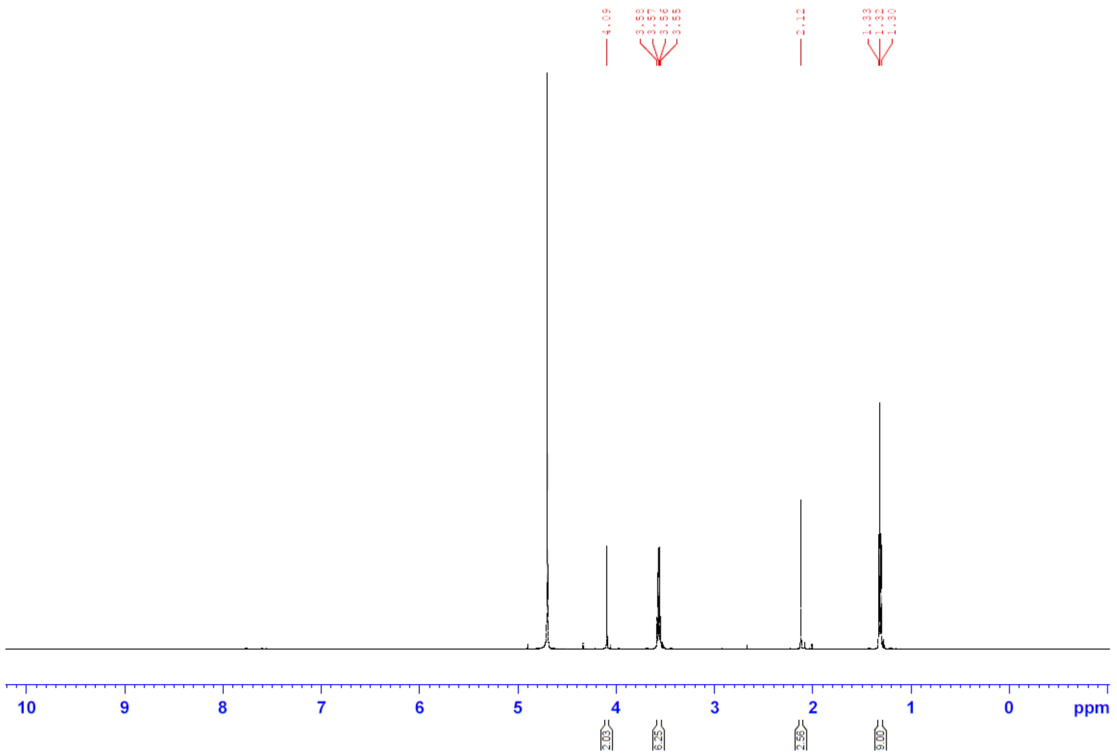

**$^{13}$C-NMR**

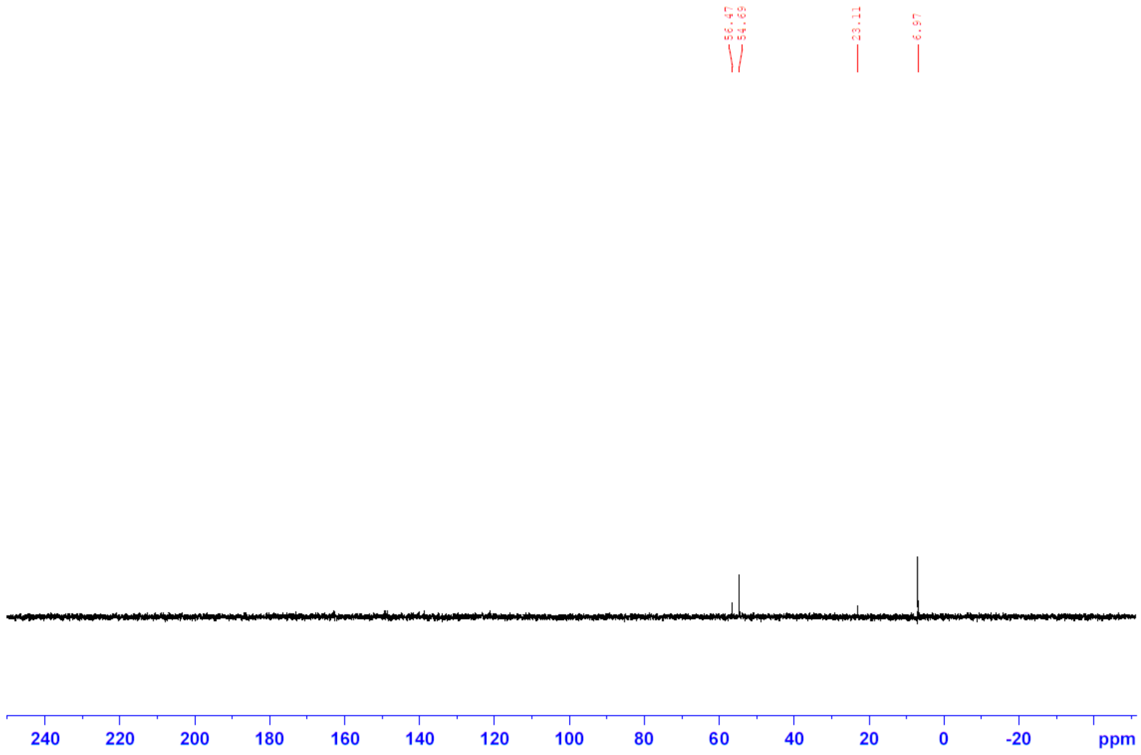

### 4. Supplementary Figures

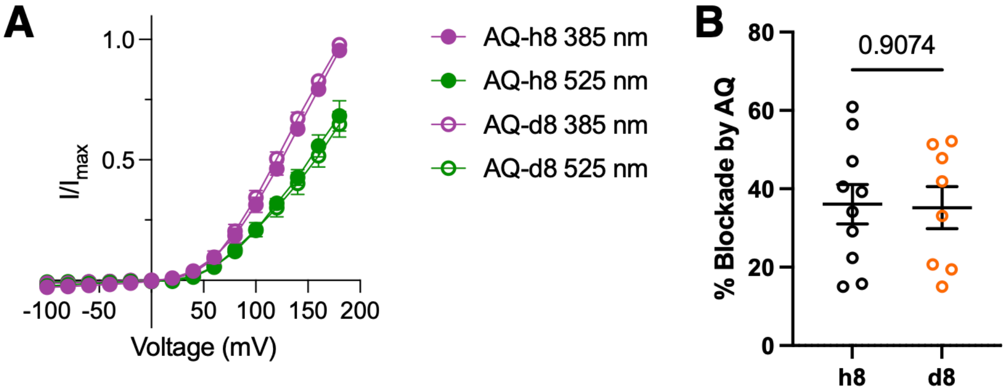

**Figure S1: BK channel blockade by AQ-h8 or AQ-d8 is comparable A)** Current-voltage relationship for BK channel activation in response to a step pulse protocol showing effect of trans-induced photo-block. **B)** Quantification of the % block for each compound calculated as 100(1-x) with x being the difference between *cis* and *trans* total current measured at +180 mV for each individual cell. P-value of unpaired t-test is reported in panel B.

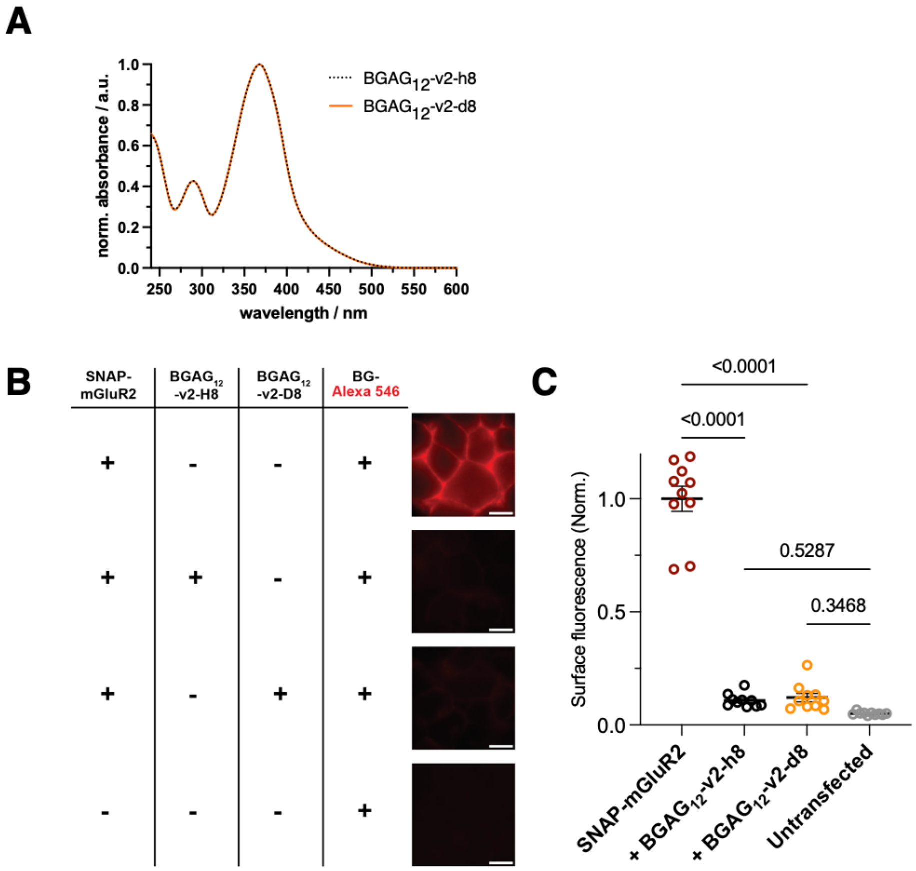

**Figure S2: BGAG photoswitching of SNAP-mGluR2. A)** UV/Vis spectra from the LCMS traces of $BGAG_{12}$-v2-h8 and $BGAG_{12}$-v2-d8. **B-C)** Labelling efficiency of the BGAG compounds. We tested the labelling efficiency of the SNAP-tag by the BGAG compounds using a pulsed-chase competition assay. P-values of One-way ANOVA with Tukey's multiple comparisons test are reported in panel C.

### 5. Supplementary Schemes

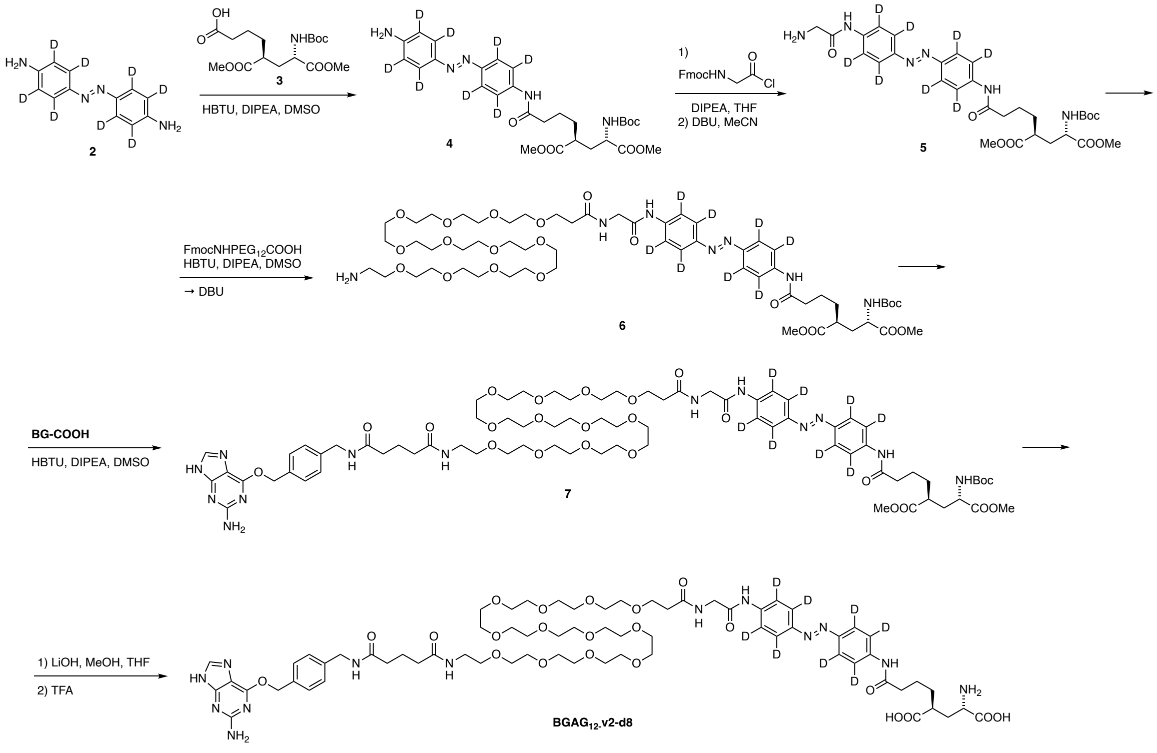

**Supporting Scheme 1: Chemical synthesis of BGAG12-v2-d8.**

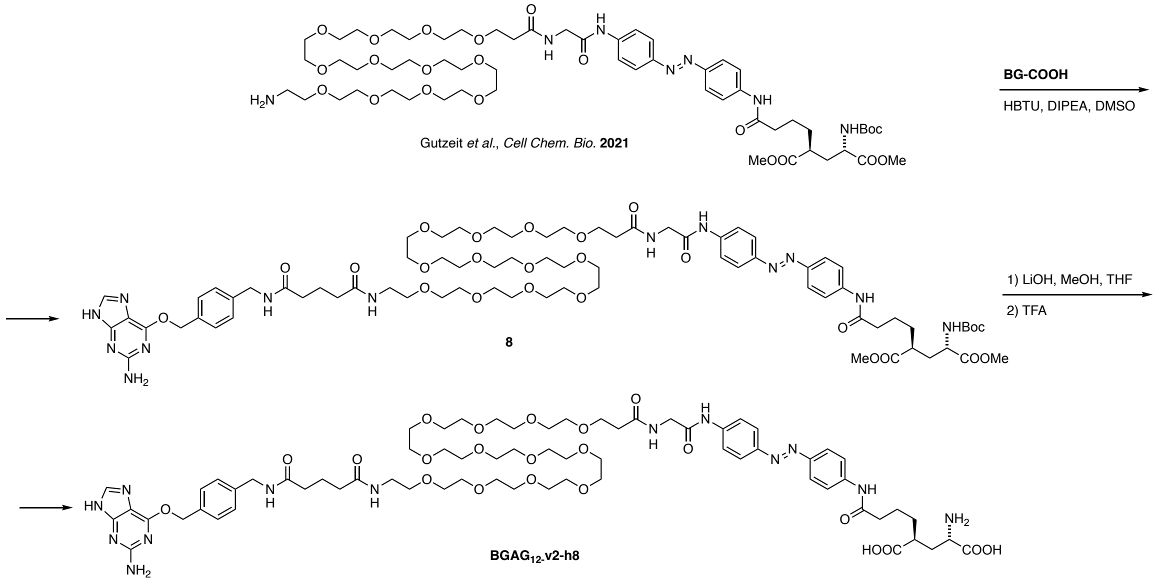

**Supporting Scheme 2: Chemical synthesis of BGAG12-v2-h8.**
